# Population genomics of the facultatively asexual duckweed *Spirodela polyrhiza*

**DOI:** 10.1101/583021

**Authors:** Eddie Ho, Magdalena Bartkowska, Stephen I. Wright, Aneil Agrawal

**Author notes:** these authors contributed equally to the work (Eddie Ho, Magdalena Bartkowska). corresponding authors contributed equally to the work.

## Abstract

- Clonal propagation allows some plant species to achieve massive population sizes quickly but also reduces the evolutionary independence of different sites in the genome.
- We examine genome-wide genetic diversity in *Spirodela polyrhiza*, a duckweed that reproduces primarily asexually.
- We find that this geographically widespread and numerically abundant species has very low levels of genetic diversity. Diversity at nonsynonymous sites relative to synonymous sites is high, suggesting that purifying selection is weak. A potential explanation for this observation is that a very low frequency of sex renders selection in effective. However, there is a pronounced decay in linkage disequilibrium over 40 kb, suggesting that though sex may be rare at the individual level it is not too infrequent at the population level. In addition, neutral diversity is affected by the physical proximity of selected sites, which would be unexpected if sex was exceedingly rare at the population level.
- The amount of genetic mixing as assessed by the decay in linkage disequilibrium is not dissimilar from selfing species such as *Arabidopsis thaliana*, yet selection appears to be much less effective in duckweed. We discuss alternative explanations for the signature of weak purifying selection.

## Introduction

The majority of eukaryotes can reproduce through outcrossing, but uniparental reproduction through asexual reproduction or self-fertilization has arisen many times independently (Vrijenhoek, 1998; Barrett, 2002; Rice, 2002; Goodwillie *et al.*, 2005; Jarne & Auld, 2006; Whitton *et al.*, 2008). Given the genetic and ecological benefits of uniparental reproduction (Fisher, 1941; Maynard-Smith, 1978; Lloyd, 1979), the rarity of highly asexual and highly selfing species in nature is a long-standing problem in evolutionary biology (Otto, 2009). Reproductive systems can have dramatic consequences on the evolution of the genome and these consequences may be key to understanding the benefits of outcrossing over uniparental reproduction. One of the major consequences of selfing and asexuality is a reduction in the effective population size, *N*_*e*_, because lower effective rates of recombination increase the strength of linked selection effects, such as background selection and selective sweeps (Golding & Strobeck, 1980; Hudson & Kaplan, 1995; Nordborg & Donnelly, 1997; Charlesworth *et al.*, 1997; Nordborg, 2000). In addition, species employing uniparental reproduction can colonize habitats with just a few individuals so population bottlenecks can further reduce local *N*_*e*_.

While Asexuality and selfing are both modes of uniparental reproduction, they differ in their effects on allelic diversity within individuals. Selfing reduces diversity by homogenizing homologous alleles within individuals. Asexuality can increase diversity within diploid individuals because there is less opportunity for recombination and segregation. This effect of alleles diverging within individuals is called the ‘Mesleson effect’ and can appear as an excess in heterozygosity (Mark Welch & Meselson, 2000; Butlin, 2002; Ament-Velásquez *et al.*, 2016). However, asexual reproduction reduces genotypic diversity within populations (Balloux *et al.*, 2003), leading to fewer distinct genotypes despite the high allelic diversity. Furthermore, in the presence of mitotic gene conversion, the effect of increased coalescence time due to clonality can be weakened or reversed, driving a reduction in nucleotide diversity (Hartfield *et al.*, 2016). Background selection will also drive strong reductions in effective population size in highly asexual lineages, an effect that can be stronger than observed for highly selfing organisms (Agrawal & Hartfield, 2016).

A decline in *N*_*e*_ is predicted to reduce neutral variation and lower the efficacy of selection such that deleterious mutations are more likely to accumulate and beneficial mutations are less likely to establish (Muller, 1932; Kimura, 1962; Heller & Maynard Smith, 1978; Charlesworth *et al.*, 1993); (Charlesworth & Charlesworth, 1997; Orr, 2000; Glémin & Ronfort, 2013; Kamran-Disfani & Agrawal, 2014). Empirical tests of these predictions increasingly rely on obtaining genomic sequences from selfing and asexual species (Wright *et al.*, 2002; Cutter, 2006; Paland & Lynch, 2006; Slotte *et al.*, 2010; Qiu *et al.*, 2011; Ness *et al.*, 2012; Brandvain *et al.*, 2013; Kamran-Disfani & Agrawal, 2014; Barraclough *et al.*, 2007; Escobar *et al.*, 2010; Tucker *et al.*, 2013; Arunkumar *et al.*, 2015; Laenen *et al.*, 2017). There is strong evidence that self-fertilization reduces neutral diversity (Schoen & Brown, 1991; Glémin *et al.*, 2006). Furthermore, highly selfing species suffer from a reduced efficacy of purifying selection as assessed from polymorphism and codon usage data, at least in some cases, though this effect is weak to non-existent when examining fixed differences between species (Glémin *et al.*, 2006; Wright *et al.*, 2002; Glémin & Galtier, 2012; Burgarella *et al.*, 2015). There may be multiple reasons for these contradicting results such as low sampling or the young age of highly selfing species (Wright *et al.*, 2002; Glémin & Galtier, 2012). In contrast, evidence for weakened purifying selection in asexual species is mostly derived from fixation data (Glémin & Muyle, 2014). Not only are there fewer studies testing the population genomic consequences of asexuality, most studies utilize mitochondrial sequence and few have used polymorphism data to test for relaxed selection (Hartfield, 2016). Given that approximately 80% of angiosperms undergo some form of asexual reproduction (Barrett, 2015; Hartfield, 2016), there is ample opportunity to study the consequences of asexuality.

Here, we analyze genomic sequences of the highly asexual aquatic duckweed, *Spirodela polyrhiza*. We are broadly interested in the population genomic consequences of the highly asexual lifestyle that characterize this duckweed. *S. polyrhiza*, the greater duckweed, is part of the Araceae family (subfamily Lemnoideae) containing 37 or 38 species of duckweed (Landolt, 1986; Les *et al.*, 2002). This aquatic angiosperm is cosmopolitan, excluding polar regions. It is thought to propagate almost exclusively through clonal reproduction (Landolt, 1986) though there are no robust estimates of sexual reproduction. Flowering was observed in 5% of samples in one survey (Hicks, 1932). We know little about *S. polyrhiza* population genomics in general. Many studies of duckweed genetics focus on species differentiation using different regions of the genome, such as AFLPs (Bog *et al.*, 2010), ISSR (Xue *et al.*, 2012) and chloroplast regions (Jordan *et al.*, 1996; Tang *et al.*, 2014; Xu *et al.*, 2015). However, most of these studies find little to no diversity within species. Crawford and Landolt (1993) examined 16 allozyme loci among 67 samples of *S. polyrhiza* isolated from Africa, Asia, Australia, Europe and North America. *S. polyrhiza* possessed a lower level of diversity than *S. intermedia* and *S. punctata*, which only had 21 and 43 isolates, respectively. They suggest that the low diversity in *S. polyrhiza* may be due to its lower frequency of seed production compared to the other two *Spirodela* species. A new population genomic study parallel to ours reports very low levels of polymorphism within *S. polyrhiza* (Xu et al., 2018).

We examine whole genome short-read sequence data from 36 *S. polyrhiza* samples. We examine patterns of diversity at different site types, how diversity varies across the genome, and patterns of linkage disequilibrium. Diversity is exceptionally low in *S. polyrhiza*, purifying selection appears to be very weak, and linkage disequilibrium extends over long distances but shows clear signs of decay. The results are consistent with sex being very low at the individual level but not too infrequent at the species level.

## Materials and Methods

### Population samples

Populations of *S. polyrhiza* were haphazardly chosen to represent the species’ genetic diversity across North America (Table S1). Of 38 samples, 26 were isolates collected from the wild across Canada and the United States. An additional 12 samples were obtained from the Rutgers University Stock Centre. Nine of the Rutgers isolates were derived from North American populations (seven from USA and two from Mexico) and the remaining three samples represent global diversity (one each from Colombia, India and France). Two of the Rutgers University samples (Texas-RU412 and Colombia-RU415) were excluded from all analyses because less than 55% of their sequenced reads mapped to the reference genome and both samples were extreme outliers in genotype distance to all other samples.

### Laboratory culturing and sequencing

All samples were established in laboratory cultures that were derived from single fronds. The cultures were grown for several months in axenic conditions at the University of Toronto. Prior to DNA extraction, fronds were collected, washed under tap water and flash frozen in liquid nitrogen. DNA was extracted from frond tissue using a modified CTAB protocol (Lutz *et al.*, 2011). Library preparation (Illumina TruSeq with PCR) and paired-end genomic sequencing were conducted at the Genome Quebec Innovation Centre at McGill University on the Illumina HiSeq2000 PE100 platform. Sequencing of the samples was conducted across three lanes generating paired-end 100 bp long reads.

### Genotyping

We used the Stampy aligner version 1.0.22 with default settings to align genomic reads to the *Spirodela polyrhiza* reference genome (Wang *et al.*, 2014). Genotyping was then performed using the Genome Analysis Toolkit (GATK) v. 2.7 GATK HaploytpeCaller using default parameters (DePristo *et al.*, 2011). The median genotype quality across all samples and sites was 29 (ranged from 16-42), and the median individual depth across all sites ranged from 5-17. We excluded two samples (Texas-RU412 and Colombia-RU415) from all analyses and called SNPs using 36 rather than 38 samples because these two (RU412 and RU415) had the lowest proportion of mapped reads, the lowest coverage and the greatest sequence divergence from all other samples.

### Hard-filtering based on sample and site quality and depth

First, we applied sample-specific filters. Sample genotypes at a site were considered missing if the sample depth was less than 5 or more than 1.5-fold the median sample depth. For variant sites, where GATK provides a genotype quality (GQ) score for individual genotypes, we excluded all genotypes with GQ score less than 20. Next, we applied a series of site-level filters. Invariant and variant sites were excluded if: 1) fewer than 20 samples had a depth between 5 and 40 reads, 2) the average sample depth exceeded 18 or was below 10, and 3) fewer than 2/3 (24) of samples passed the sample-specific filters. Filtering for high depth was performed to avoid regions with paralogous read mapping; SNPs at sites with high average sample depth were more likely to have fixed heterozygote sites (where all samples were heterozygous). Variant sites were retained if at least 20 samples had a genotype quality (GQ) score at least 20 (which corresponds to a genotyping accuracy of 99) and if the mapping quality (MQ) of the site was at least 90. Finally, we filtered entire regions of the genome. In particular, we removed sites that were identified as transposable elements or highly repetitive along with 100 bp on either side of the repetitive element. We used the repeat-masked assembly available on JGI (Spolyrhiza_290_v2.repeatmasked_assembly_v1.gff3), which masked retroelements, as well as RepeatModeler (v. 1.0.8) coupled with RepeatMasker (v. 4.0.5) to identify transposable elements and highly repetitive regions. Indels were removed along with 5 bp on either side of the site; for a deletion we also removed all sites spanning the length of the deletion. 20 kb windows were removed if fewer than 40% of sites within the window failed to pass all other filters. This filter eliminated regions that tended to have abnormally high number of poorly mapped reads and tended to have clusters of highly variable sites near poorly assembled regions of the genome. For gene-level analyses, we also removed 1595 genes (out of the 19623 annotated genes) that had sites where the average sample depth was greater than 18 reads, to further filter out paralogous genes.

### Genetic distance

We calculated pairwise genetic distance using two methods. The pairwise ‘genotypic distance’ was calculated by summing the number of sites between two samples that differed in genotype. Pairs of sites that are homozygous for different alleles (homozygous differences) are weighted twice as much as pair of sites where one sample is homozygous and the other samples is heterozygous (heterozygous differences) (Table S2). The pairwise ‘allelic distance’ is the probability that a randomly selected allele from two samples at a given site will be different. The two distance metrics differ only in how they score the distance between samples that are both heterozygotes: the genotypic distance is zero but the allelic distance is ½, as it is for homozygote vs. heterozygote comparison (Table S3). Comparisons of the two metrics are useful when there are high rates of asexual reproduction as there may be low genotypic diversity despite a retention of allelic variation. We used these pairwise genetic distances to construct neighbour joining trees using the Ape package (Paradis *et al.*, 2004; Paradis, 2011) in R (R Core Team, N, 2016).

### Grouping samples into genets

Because *S. polyrhiza* reproduces asexually, it is very likely that samples from the same or nearby pond may be clones (i.e., descended from a common ancestor via only clonal reproduction, possibly over many generations). For most analyses, we grouped samples that were highly genotypically similar into one genet that was genetically distinct from other genets. Samples were grouped into genets as follows. For each pair of samples, we calculated the number of sites where one sample is heterozygous and the other is homozygous (“heterozygous differences”) and the number of sites where one samples is homozygous for one allele and the other is homozygous for the other allele (“homozygous differences”). Clonal samples will not necessarily be completely identical because of genotyping error (though bioinformatics filtering should minimize this) and because of mutation and gene conversion. Nonetheless, clonal samples should be very similar. If two samples are related exclusively through clonal propagation, heterozygous differences can occur simply due to a point mutation in one of the lineages. With clonal reproduction, homozygous differences will be even rarer because they require two rare events, e.g., two point mutations occurring at the same site or two lineages to be separated by both mutation and a gene conversion event. Based on an inspection of the data (see Results and Discussion), we clustered samples together if each pair within the cluster had ≤ 0.01% of sites with homozygous differences and ≤ 2% of sites with heterozygous differences, as we found these thresholds to form very distinct groups. Using this threshold, we found nine genotypes composed of multiple samples and three genotypes each represented by a single sample (Table S5). For each of the 12 genotypes (hereafter, genets) we created a ‘consensus genotype’ whereby the genotype of each site was randomly chosen among the samples that form the group. Pairwise genotypic distance between samples within genets was, on average, 35 times smaller than genotypic distances between samples from different genets (Table S4). We confirmed our grouping using k-means clustering (Adengenet package in R; Jombart & Ahmed, 2011). This procedure consists of running successive K-means with an increasing number of clusters (k), after transforming the data using a principal component analysis (PCA).

These analyses found that one genet (consisting of RU448 and RU99) was very divergent from the other genotypes and possessed much higher heterozygosity (below). For subsequent analyses, we excluded this genet and only included the remaining 11 genets for all subsequent analyses.

### Estimating genetic diversity

Gene annotations for *S. polyrhiza* were obtained from the Joint Genome Institute’s Phytozome (https://phytozome.jgi.doe.gov/pz/portal.html#!info?alias=Org_Spolyrhiza). We used this annotation to obtain genetic diversity estimates for different site types. After filtering, 15310 of the 19623 annotated genes in the *S. polyrhiza* reference genome remained. We categorized these genes based on their level of expression in fronds and turions (characterized by Wang *et al.* 2014, FKPM values provided by Joachim Messing, Rutgers University) and homology to three monocot species (*Sorghum bicolor, Zea mays*, and *Oryza sativa*). Of these, 1058 genes had no detectable level of expression (FPKM was 0 in both fronds and turions), leaving 14252 genes in the expression groups. We separated these remaining genes into four categories of expression, each with 3562 genes (low, mid-low, mid-high and high).

We further split these genes based on three levels of evolutionary constraint by using Blastx to assess sequence similarity between *S. polyrhiza* genes and three other species (*Sorghum bicolor, Zea mays, Oryza sativa*). We found genes with blast hits to all three species. For each *S. polyrhiza* gene we summed the bit scores for each of the three blast hits and categorized genes into 3 levels of constraint (high, mid and low).

### Linkage disequilibrium

The pattern of LD decay across all genotypes should reflect historical recombination events between the disparate genotypes. We calculated LD as the covariance across diploid genotypes using the script provided by Rogers and Huff (2009) to estimate the covariance between two vectors of genotypes. We modified this script to calculate LD across all loci where the minor allele frequency was greater than 0.05. Within each scaffold we calculated LD between pairs of sites up to 500 kb. We also estimated the average *r*^2^ between pairs of scaffolds to establish genome-wide background levels of LD. For this, we randomly sampled one site for each of 32 scaffolds and calculated all pairwise *r*^2^ among these 32 sites. We repeated this 10000 times to get an estimate of *r*^2^ across scaffolds; we got similar results if we only repeated the sampling 500 or 1000 times.

### Recombination hotspots

We used the ‘rhomap’ function within LDhat (McVean *et al.*, 2004) to estimate *ρ* = 4*N*_*e*_*r* for each scaffold separately, where *r* is the recombination rate per generation. We ran the program for 9,900,000 iterations sampling every 1000 iterations after a burn-in of 100,000 iterations with a block penalty of 5. To search for putative recombination hotspots, we used the output of LDhat to calculate the average recombination rate of 1kb non-overlapping window within each scaffold. We then putatively define recombination hotspots as windows that have 10 times the recombination rate of the scaffold average; adjacent windows that pass this threshold were merged into one putative hotspot.

To examine whether the putative recombination hotspots possessed unique characteristics relative to the rest of the genome we sampled a set of ‘control regions in the genome. We randomly sampled 131 regions of the genome with the same length as the putative hotspots requiring that each region had <= 5% missing nucleotides in the reference, at least one SNP and did not overlap with existing hotspots or other control regions. We then examined differences in the GC content, proportion of bases overlapping coding regions and genetic diversity between our hotspots and the control regions using logistic regressions in R (R Core Team 2016). The logistic model we used was: Region type (i.e. hotspot or control) ∼ GC content + genetic diversity + coding sequence overlap.

## Results and Discussion

### Heterozygosity and clonal genotypic structure

After variant calling and filtering, we obtained 79,166,349 invariant and 417,884 variant sites among the 36 *S. polyrhiza* samples, which resulted in an average observed heterozygosity of 0.000636 per site (Table S5). Grouping the samples into 12 distinct genets (i.e., 12 genotypes that are not clonal relatives of one another) resulted in a similar average observed heterozygosity of 0.000694 (Table S4). However, one genet (made up of samples RU448 and RU99) had approximately three times higher heterozygosity compared to the average of the other 11 genotypes (Table S4). This genotype was also highly divergent from the others (below) and over-contributed to the singleton category in the allele frequency spectrum. For these reasons, this genotype was excluded from most analyses below (unless stated otherwise). For the remaining 11 genets, we had 71,949,140 invariant sites and 142,106 variant sites and the average observed heterozygosity was 0.000588. We calculated F_IS_ = 1 – H_obs_ / H_exp_ among the 11 genets at each site and found the average F_IS_ to be 0.044, where H_obs_ and H_exp_ are the observed and expected heterozygosity at a site, respectively (Figure 1).

**Figure 1.**
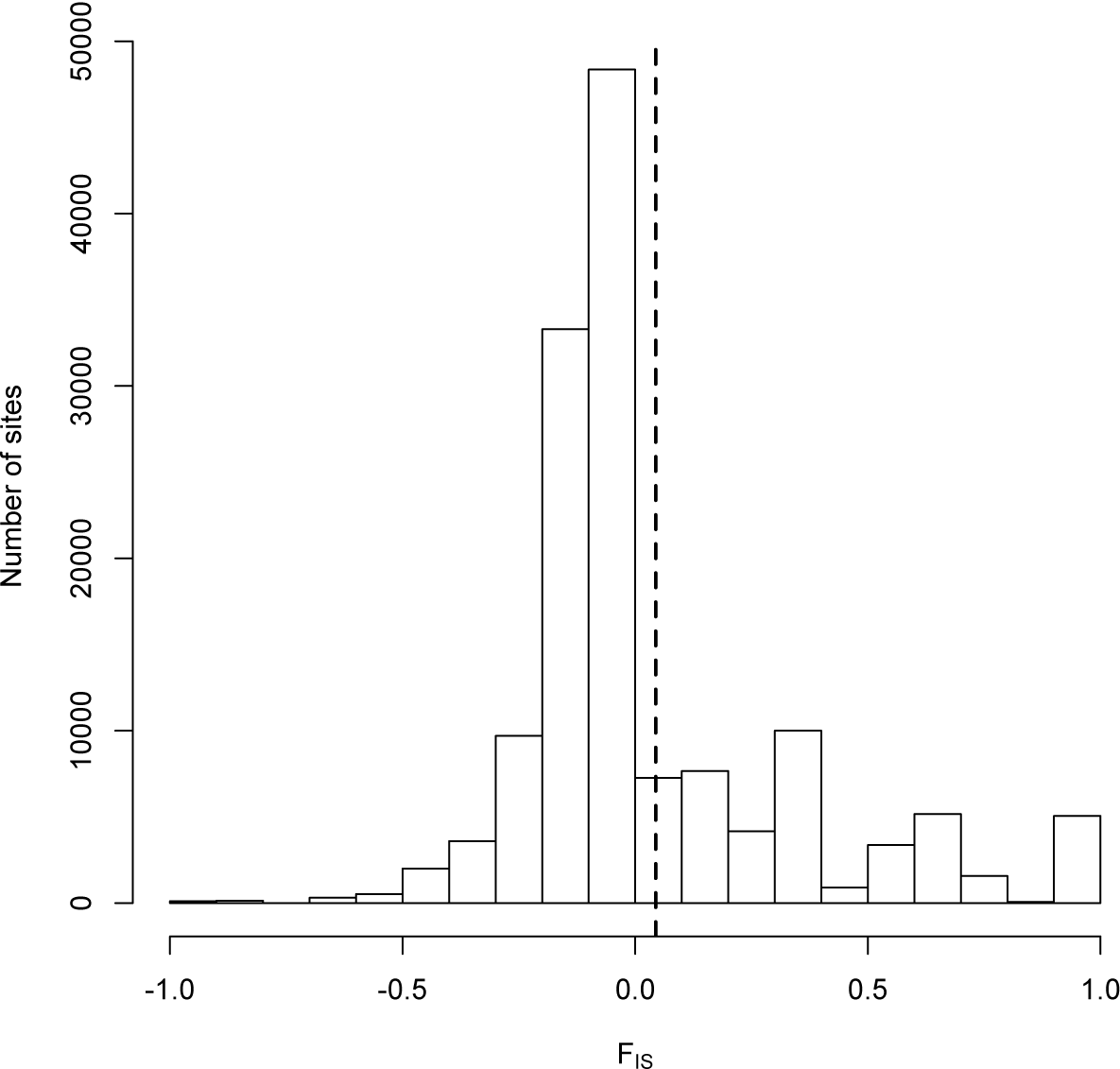
Distribution of F_IS_ for 11 *S. polyrhiza* genets.

### Genetic distance

We constructed neighbour joining trees based on their pairwise genotypic and allelic distance among the 36 *S. polyrhiza* samples (Figure 2). Both trees revealed consistent clusters of genetically similar samples and the one pair of samples (RU99 and RU448) that was highly divergent. The 36 samples clustered into 12 groups based on the number of homozygous and heterozygous differences between pairs of samples (Table S4, Fig. S1). Individuals found in the same genotype group generally showed strong geographic associations; for example, all samples from Oklahoma belonged to the same genotype group (samples labelled CC and RC). Our most extensively sampled population from a large pond in Toronto (GP) had five samples from a single genotype group (GP8-1, GP10-3, GP6-5, GP4-2, and GP2-3), while a sixth sample (GP4-4) formed a separate genotype group with a sample from a nearby pond. On the other hand, there is no significant correlation between genetic and geographic distance among the 12 genets (genotypic distance: Pearson’s *r*=0.092, *p*=0. 16; allelic distance: *r*=0.041, *p*=0.54) Three samples (RD24, BC RR2_1 and RU195) were the sole representative of their genet while all other genotypes had at least two samples (Table S4). We confirmed the groupings into distinct genets using k-means clustering. The model with 13 genets had the lowest BIC score (BIC= 305.7751), but that with 12 groups was the next best fit (BIC= 307.1549). Clustering with k = 13 assigned the two Nova Scotia samples (HFA10, HFB11) to separate groups. However, based on the low pairwise genotypic and allelic distance between these two samples, we decided to represent the two samples as one genet in downstream analyses.

**Figure 2.**
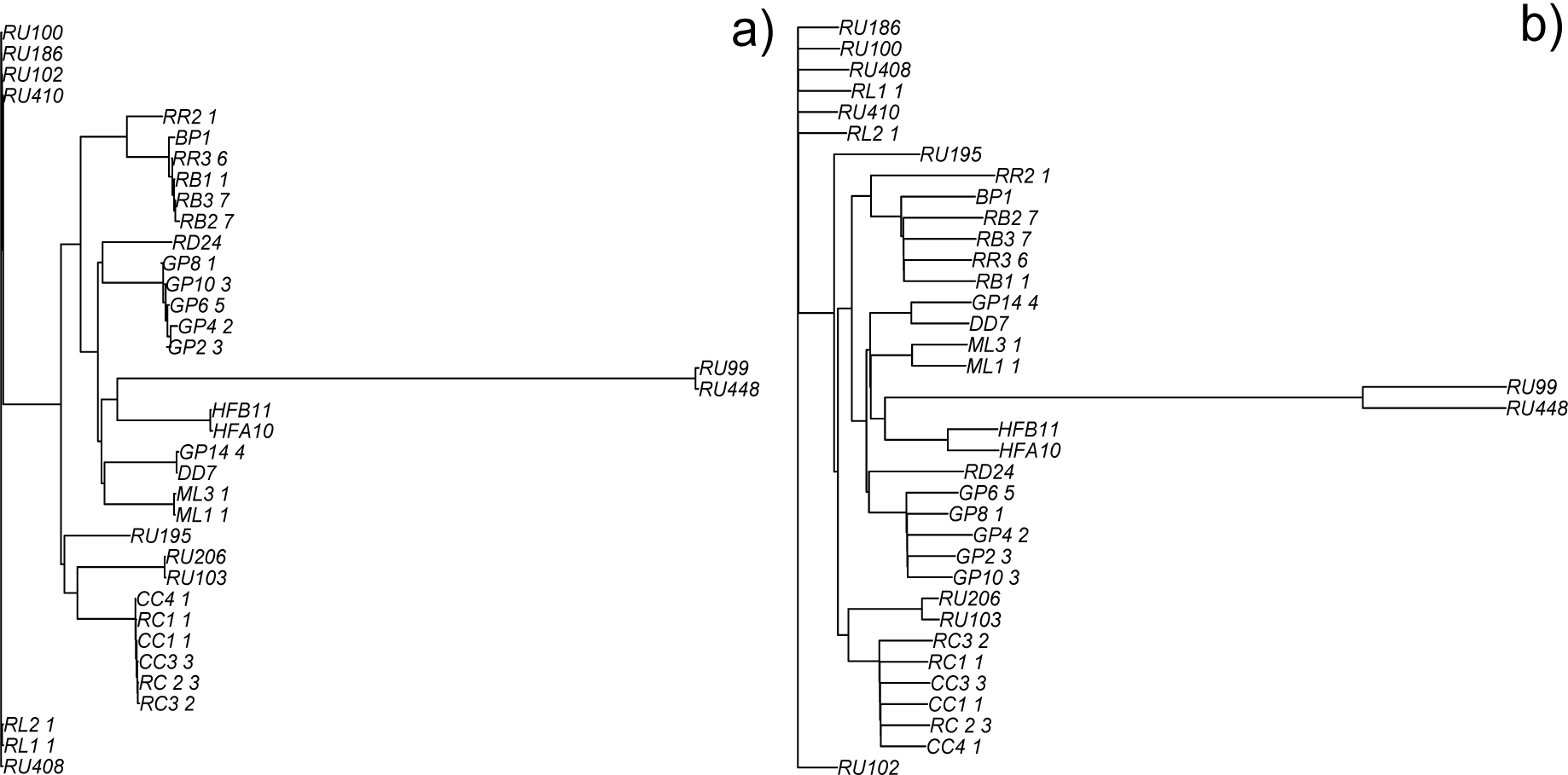
Neighbour joining trees for 36 *S. polyrhiza* samples constructed using pairwise (a) genotypic and (b) allelic distances.

The unusual genet consisting of RU99 and RU448 was divergent in genotypic and allelic distances suggesting that was very different from the other genets. To investigate this further we examined sites where a focal genet possesses an alternate allele (i.e. heterozygote or homozygous for the alternate allele) but all other genotypes are homozygous for the reference allele. We found that the {RU99, RU448} genet possess 223652 sites where it is the only genet with the alternate allele, while the other 11 genotypes possess 7542 of these sites, on average (Table S4). Due to the strong differentiation of the {RU99, RU448} genet from the others and its abnormally high levels of heterozygosity (above), we removed it from downstream analyses.

### Genetic diversity

Among the 11 *S. polyrhiza* genets, genetic diversity across all sites was very low (*π* = 0.000542) compared to other plants (Chen *et al* 2017). We estimated *π*_*s*_ to be 0.000463 at 4-fold degenerate (synonymous) sites and *π*_*n*_ to be 0.000229 at 0-fold (non-synonymous sites) resulting in *π*_*n*_ /*π*_*s*_ = 0.495 (Table 1), suggesting a relatively low level of selective constraint in *Spirodela* on amino acid mutations. This ratio is considerably higher than other plant species estimates to date (Chen *et al* 2017). Diversity at 2- or 3-fold degenerate sites was 0.000351 and as expected falls between diversity at 0-fold and 4-fold sites. Diversity at intergenic sites was 0.000732, which was higher than *π*_*n*_. Values were similar for Watterson’s estimate of diversity (*θ*_*w*_) and *π* resulting in Tajima’s *D* values close to zero for both synonymous and intergenic sites (Table 1). Tajima’s *D* is slightly more negative at nonsynonymous sites, consistent with weak purifying selection.

**Table 1.**
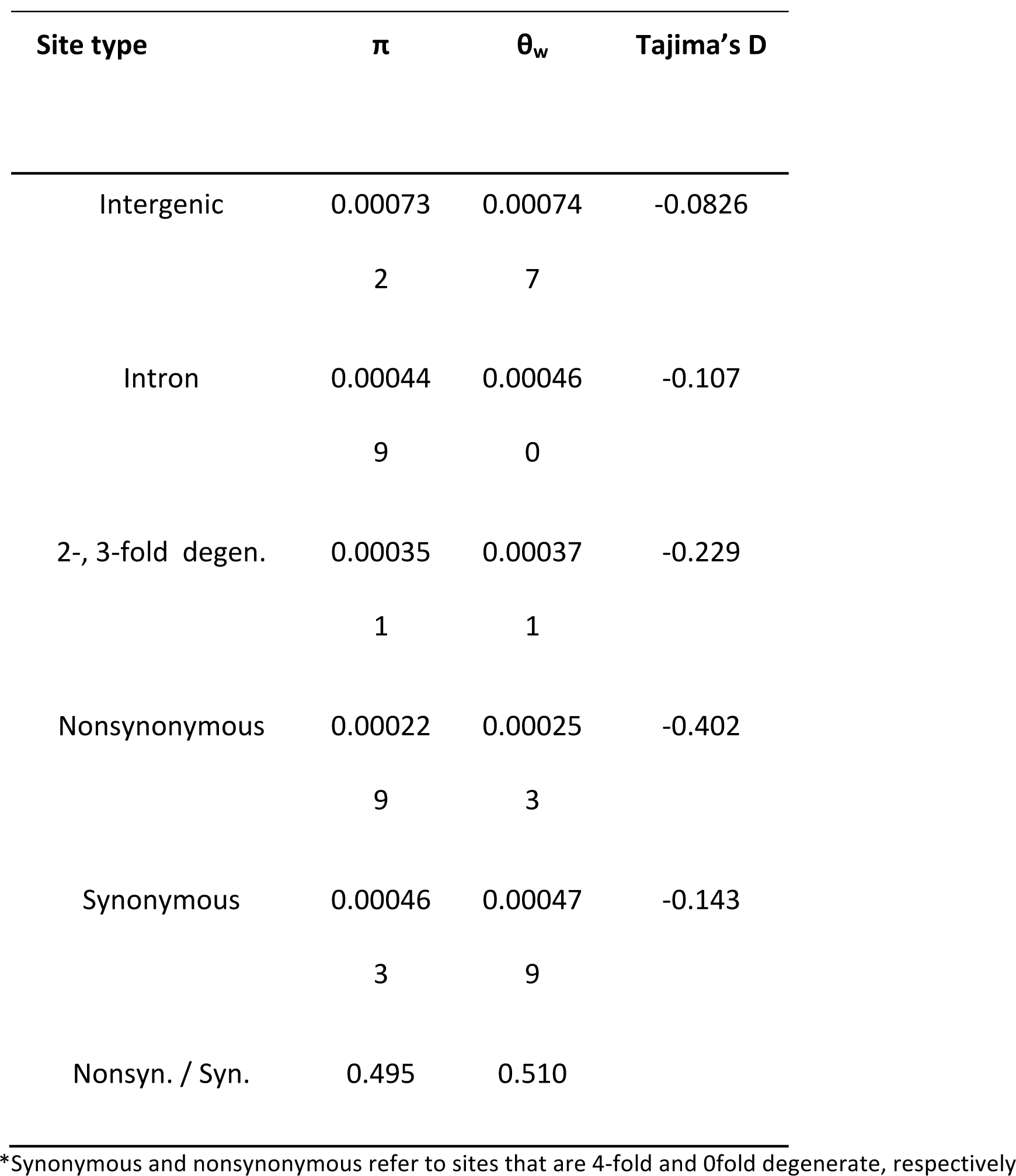
Genetic diversity and Tajima’s D estimates.

The relatively high value of *π*_*n*_ /*π*_*s*_ could reflect a reduced genome-wide efficacy of natural selection (*N*_*e*_*s*) due to low rates of sexual reproduction causing low *N*_*e*_ but may also reflect weaker selection (i.e., low *s*) on many genes, perhaps due to a simplified morphology and life cycle since divergence from monocot ancestors. Under the latter hypothesis, the elevated *π*_*n*_ /*π*_*s*_ may only occur in genes with little to no expression. Genes with lower expression had higher diversity at both synonymous and non-synonymous sites (Figure 3, Table S6). However, the reduction in diversity with increasing expression levels was proportionally faster for *π*_*n*_ than for *π*_*s*_, which resulted in *π*_*n*_ /*π*_*s*_ values decreasing with increasing expression (e.g. *π*_*n*_ /*π*_*s*_ for low expression and high expression genes was 0.557 and 0.346, respectively). These results are consistent with purifying selection being stronger at genes with higher expression levels, consistent with patterns observed in other species (Carneiro *et al.*, 2012); (Paape *et al.*, 2013; Williamson *et al.*, 2014). Very high levels of *π*_*n*_ /*π*_*s*_ in genes with low or no expression may indicate that some fraction of these genes are not functionally important. However, even genes in the more highly expressed categories show relatively high *π*_*n*_ /*π*_*s*_ compared to other plants (Chen *et al.*, 2017), suggesting a genome-wide signal of low selection efficacy in *S. polyrhiza*.

**Figure 3.**
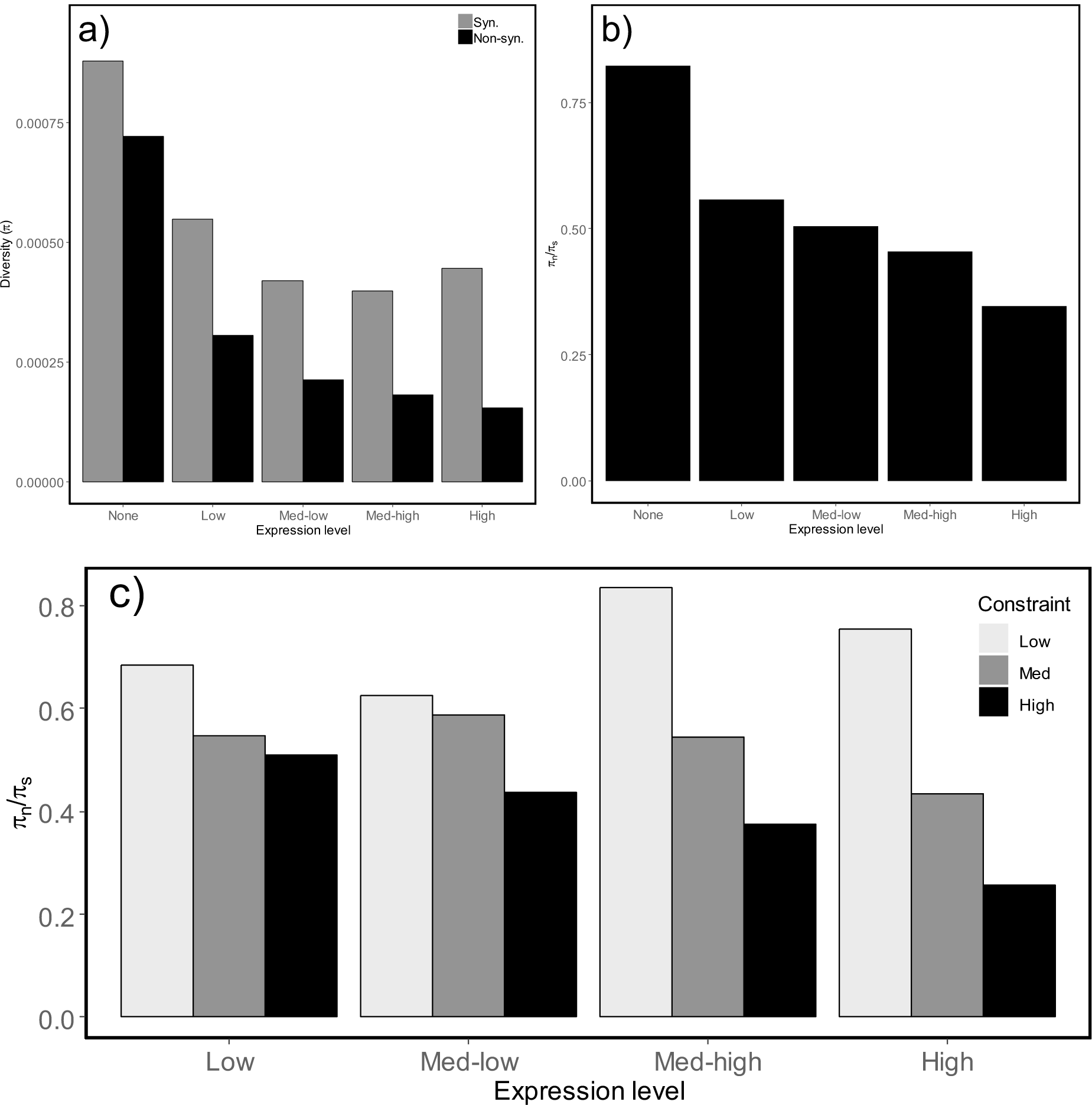
(a) π_s_ and π_n_ for *S. polyrhiza* at genes with varying expression levels. (b) π_n_ / π_s_ in *S. polyrhiza* genes that have different expression levels within *S. polyrhiza* tissue. (c) π_n_ / π_s_ in *S. polyrhiza* genes that have different expression levels within *S. polyrhiza* tissue. Within each expression level category, genes are separated into low, mid and high evolutionary constraint based on their divergence to homologous genes in *Sorghum bicolor, Zea mays, Oryza sativa*. *S. polyrrhiza* consists of 11 genets.

To further explore variation across genes in *π*_*n*_ /*π*_*s*_, we categorized *S. polyrhiza* genes within each expression level category based on their evolutionary constraint, based on their *blastx* divergence from *Sorghum bicolor, Zea mays, Oryza sativa*. We observed that within each expression category, genes that are more conserved had lower *π*_*n*_ /*π*_*s*_ values (Figure 3, Table S7-S10). Nevertheless, *π*_*n*_ /*π*_*s*_ values are high in all expression/constraint categories. Only in the most highly expressed and highly constrained genes does *π*_*n*_ /*π*_*s*_ approach values typically observed in outcrossing plants (Chen *et al.*, 2017).

There are two patterns in synonymous diversity that are somewhat unexpected. Diversity at synonymous sites is low relative to intergenic sites, and it is also elevated in genes that are weakly expressed compared to those in high expression categories. There are two possible explanations for these patterns; synonymous sites may themselves be under purifying selection or they may be subject to the effects of background selection (or other forms of linked selection) from neighboring selected sites. If background selection is acting, we would predict that synonymous diversity should be reduced in regions with a higher density of functional sites. To test this, we examined the relationship between diversity across 50 kb windows for different site types using a linear model that includes coding site (CDS) density, GC content, and the CpG/GpC ratio, which has been used as an indicator of the level of DNA methylation (i.e. higher CpG/GpC implies less methylation (Hellsten *et al.*, 2013). As shown in Table 2, CDS density negatively affects diversity at all site types. The effect on synonymous sites is marginally nonsignificant but in the same direction as for other site types. The lack of significance may arise from the large number of windows lacking any synonymous SNPs, likely violating the assumptions of the model. Re-analyzing the data using only windows containing at least one SNP of each site type, reveals that the effect of CDS density is highly significant on all site types, including synonymous sites (Table S11). These results are consistent with background selection reducing diversity in regions with more sites under constraint. The CpG/GpC ratio has a significantly negative effect on diversity, most notably for intergenic sites, consistent with the idea that regions with low methylation (i.e., high CpG/GpC) have lower mutation rates.

**Table 2.** Coefficients from linear model of diversity (*π*) of different site types (per 50 kb window) as function of coding site (CDS) density, % GC, and the CpG/GpC ratio.

| Site type | Variable | Estimate | t | P-value |
| --- | --- | --- | --- | --- |
| Intergenic | CDS density | -0.00061 | -2.17 | 0.030 |
|  | % GC, | -0.00038 | -1.18 | 0.237 |
|  | CpG:GpC | -0.00063 | -4.71 | < 0.0001 |
| Intronic | CDS density | -0.00096 | -4.52 | < 0.0001 |
|  | % GC, | 0.00017 | 0.72 | 0.474 |
|  | CpG:GpC | -0.00021 | -2.06 | 0.039 |
| 0 fold | CDS density | -0.00102 | -5.40 | < 0.0001 |
|  | % GC, | 0.00044 | 2.02 | 0.043 |
|  | CpG:GpC | -0.00006 | -0.64 | 0.523 |
| 4 fold | CDS density | -0.00056 | -1.59 | 0.111 |
|  | % GC, | -0.00008 | -0.19 | 0.848 |
|  | CpG:GpC | 0.00007 | 0.44 | 0.661 |

### Linkage disequilibrium and recombination heterogeneity

Levels of linkage disequilibrium (LD) among the genets are affected by the amount of recombination that has occurred in the past. Among the 11 *S. polyrhiza* genets, we observed that sites within 1 to 20 bp of each other had an average *r*^2^ of 0.57 that decays to approximately 0.23 after a distance of 20 kb (Figs. 4a, b), after which there is a slower LD decay that continues to 100 kb. Between-scaffold LD is slightly but significantly lower (0.13) than LD at 100 kb (0.15), implying small residual LD at very large distances. This pattern of LD decay is comparable to that seen in the highly self-fertilizing species *Arabidopsis thaliana* (10-50kb, depending on sampling; (Nordborg *et al.*, 2005; Kim *et al.*, 2007) and *Medicago truncatula* (Branca *et al.*, 2011), whereas outcrossing populations often show much more rapid LD decay over several hundred base pairs (Foxe *et al.*, 2009; Mackay *et al.*, 2012).

**Figure 4.**
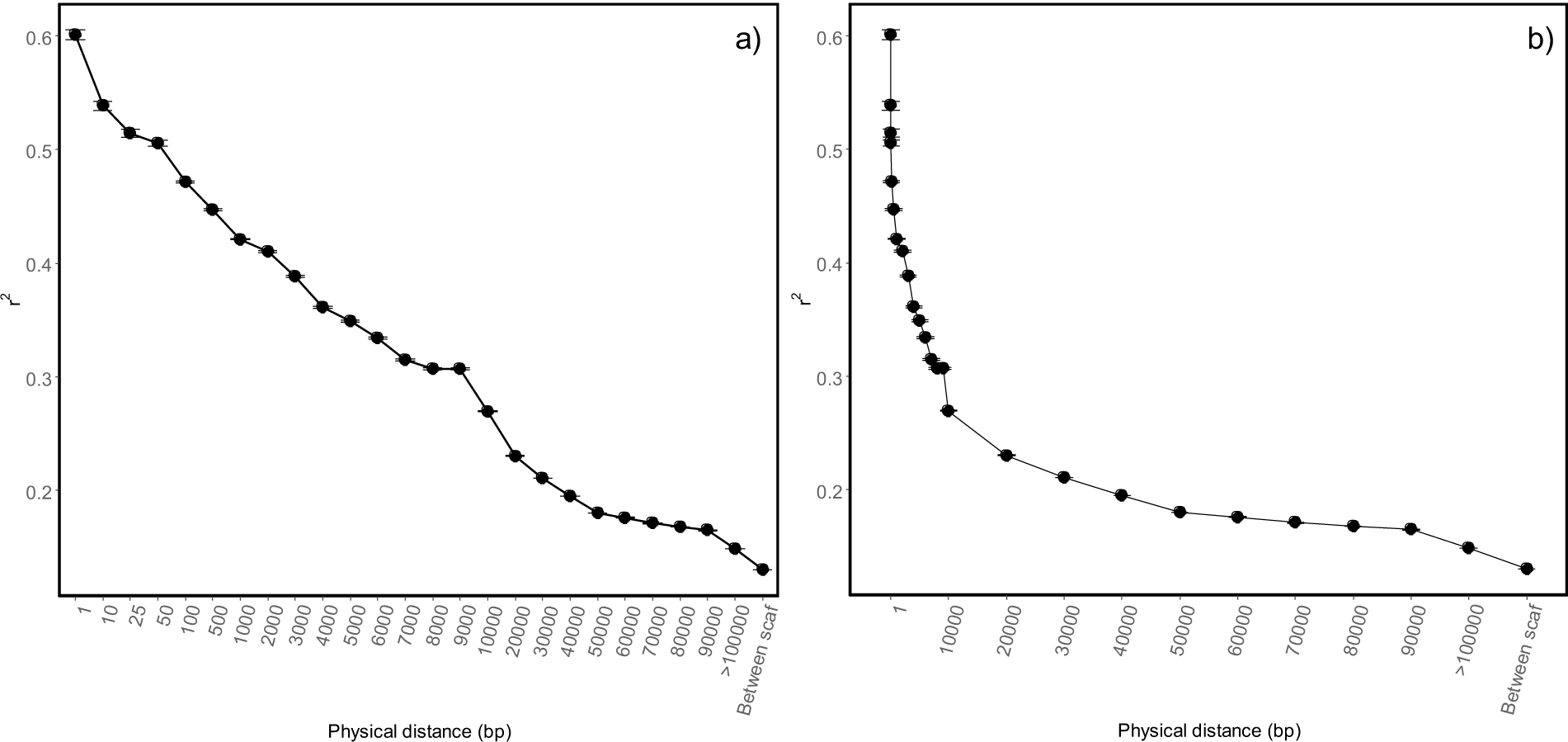
(a) Linkage disequilibrium among 11 genets of *S. polyrhiza*, measured as mean *r*^2^, decaying with distance (bp) between pairs of sites. Each point contains *r*^2^ values binned by physical distance. For example, bins contain pairs of loci that are 1-10, 10-25, 50-100, …, etc. bp apart. The last point indicates the average *r*^2^ between scaffolds. (b) Same as (a) but with the distance shown on a linear scale.

On average across our 50 kb windows, our estimate of the population recombination rate *ρ* from LDhat is 0.00051. The ratio of *ρ*/*θ* is thus approximately 1, implying an effectively comparable rate of recombination and mutation. This ratio is considerably higher than that estimated in highly selfing species (Nordborg *et al.*, 2005; Branca *et al.*, 2011), while it is on the same order as outcrossing species (Wright *et al.*, 2003; Langley *et al.*, 2012). Since *Spirodela* is predominantly asexual, this high ratio of *ρ*/*θ* is somewhat surprising. There are several possibilities that might explain this discrepancy. First, the very low level of neutral diversity may reflect a very low mutation rate, potentially due to the low numbers of cell divisions per generation compared to other vascular plants. Recent estimates of mutation rate in this species are consistent with this possibility (Xu *et al.*, 2018; Sandler et al. in prep). Second, in partially asexual species, mitotic gene conversion and mitotic crossing over play a major role in reducing linkage disequilibrium in facultatively sexual organisms, but would not do so in selfers (Hartfield *et al.*, 2018). Thus, the combination of high ρ/θ and very low diversity is not inconsistent with predictions of a highly clonal organisms experiencing mitotic recombination and/or a low rate of per-base mutation.

Using LDhat, we identified 131 putative recombination hotspots in *S. polyrhiza* that possessed a recombination rate ten times higher than the average of the scaffold they belong to (Table S12). Compared to some species (e.g. mammals), these hotspots are weak in strength and very few in number. One possible explanation for their apparent rarity is that low levels of polymorphism are causing low power to detect hotspots. Indeed, hotspot presence was significantly correlated with levels of nucleotide diversity (*π*: Pearson’s *r* = 0.28, *p*<0.0001,*θ*_*W*_: *r* = 0.31, *p*<0.0001;). To test whether low SNP density limits our detection power, we randomly masked SNPs from our data such that each 2 kb had a maximum of 5 SNPs and then re-ran LDhat and resampled appropriate control regions. In this SNP-masked dataset, we found only 50 putative hotspots which recovered ∼36% of the putative hotspots in the full dataset; two of the 50 putative hotspots were not found in the full dataset. This suggests that there is bias in the unmasked dataset for detecting hotspots in SNP-dense regions.

To look more broadly at recombination rate heterogeneity, we estimated average *ρ* in 50 kb windows across the genome, and tested for correlations with several genomic features. Previous work in plants suggest that recombination rates are elevated upstream of coding regions, and are associated with demethylated promoter regions. For these reasons, we examined the effects of coding sequence density and the CpG/GpC ratio. *ρ* is significantly positively correlated with coding sequence density (Pearson’s *r=*0.20, *p*<0.0001), GC content (*r=*0.13, *p*<0.0001), CpG/GpC (*r=*0.16, *p*<0.0001) and *π* (*r* = 0.08, p = 0.005). Since these variables are likely to be non-independent of each other, we also ran the linear model *(ρ ∼ GC content+CpG/GpC+coding sequence density+ π*) (Table 3). From this analysis, both coding sequence density and CpG/GpC positively affect *ρ*, while GC content now has a negative effect. The effects of coding density and CpG/GpC are consistent with recombination being biased towards open chromatin, as seen in other plants (Hellsten *et al.*, 2013; Rodgers-Melnick *et al.*, 2016).

**Table 3.** Linear model examining variation in estimates of *ρ* for 50 kb windows as a function of window characters: CDS density, % GC, the CpG/GpC ratio and diversity (*π*). Statistical model: lm(rho ∼ CDS density + % GC+ CpG/GpC ratio + *π*)

|  | Estimate | Standard<br>error | *t* | *p* |
| --- | --- | --- | --- | --- |
| Intercept | 0.5236 | 0.1127 | 4.645 | < 0.0001 |
| CDS density | 0.8997 | 0.1833 | 4.909 | < 0.0001 |
| % GC | -0.8643 | 0.3647 | -2.37 | 0.0179 |
| CpG:GpC | 0.2687 | 0.0989 | 2.716 | 0.0067 |
| $\pi$ | -30.0521 | 20.2418 | -1.485 | 0.1379 |

## Conclusion

The observation of low diversity could be explained by either low mutation rate or a low *N*_*e*_. Direct estimates of the mutation rate indicate that the mutation rate is very low (i.e., one to two orders of magnitude lower than *Arabidopsis*; Xu *et al.*, 2018, Sandler et al *in prep*). Based on this low estimate, Xu *et al* (2018) inferred that *N*_*e*_ is quite large (∼10^6^); our own estimate of the mutation rate (Sandler et al. in prep) and diversity is broadly consistent with a large *N*_*e*_.

At the outset, we had expected to find *N*_*e*_ to be low because this plant produces primarily by cloning so the effects of linked selection could be large (Agrawal and Hartfield 2016). However, background selection will not reduce *N*_*e*_ as much as it would if the mutation rate was higher. Nonetheless, the effects of linked selection are important in this system as *N*_*e*_ is likely much smaller than *N*. While no direct estimate of the census population size *N* exists, we expect it is massive. There are ∼6 million ponds across Canada and the USA [flyways.us] so it is not unreasonable to speculate that census population size exceeds 10^9^.

Our observation that *ρ*/*θ* is close to 1 implies that the effective recombination is close to the mutation rate. From this, we can infer the effective recombination rate in this species is several orders of magnitude lower than it is in outcrossing species such as *Drosophila* which also have ρ/θ values close to 1 but also have much higher mutation rates than in *Spirodela*. Presumably, this low effective recombination rate occurs because of the low rate of sex. If the recombination rate per meiosis is 10^−8^ (comparable to other species), but the effective recombination rate is 10^−10^ (i.e., equal to the mutation rate, see Xu et al. 2018), then we infer the rate of sex is 10^−10^/10^−8^ = 10^−2^. This simplistic calculation ignores the potential importance of mitotic gene conversion (which contributes to the inferred estimate of *ρ*), so the true rate of sex may be substantially lower. Thus, low mutation rate and low rates of sexual reproduction are likely contributing to our patterns of diversity and linkage disequilibrium.

Though we infer the rate of sex is low at the individual level, we see evidence of the effects of recombination at the species level. First, linkage disequilibrium declines with distance. The decline in LD is not dissimilar to that observed in selfing plants such as *Arabidopsis thaliana* (Nordborg *et al.*, 2005; Kim *et al.*, 2007) and *Medicago truncatula* (Branca *et al.*, 2011) that outcross at a low rate. Second, *π*_*s*_ is lower in gene dense regions, a pattern expected if recombination localizes background selection effects to tightly linked regions of the genome.

The most perplexing observation is the high *π*_*n*_/*π*_*s*_ relative to other species, given that the effective population size is estimated to be large (Xu et al. 2018). The simplest explanation is that selection (*s*) tends to be weak in *S. polyrhiza*. This could be because of relaxed selection on many genes due to the diminutive form and lifestyle relative to other angiosperms. Alternatively, selection may not be realized much of the time because local populations often consist of a single clone so there is not competitive selection among genotypes. Finally, the high *π*_*n*_/*π*_*s*_ might be explained by non-equilibrium conditions. It has been pointed out that neutral diversity takes longer to build up to its equilibrium levels than selected diversity, which can result in a transiently elevated *π*_*n*_/*π*_*s*_ (Simons *et al.*, 2014; Brandvain & Wright, 2016). Our estimates of Tajima’s D are negative, which could reflect recovery from bottlenecks. Disentangling the reasons for the high *π*_*n*_/*π*_*s*_ remains a challenge for future work.

## Acknowledgements

This work was supported by the Natural Sciences and Engineering Research Council of Canada (AFA and SIW). We thank Jade Lavallee, Victor Mollov, and Niroshini Epitawalage for help with plant growth and extractions and Adrian Platts for bioinformatics assistance.

