## Supplemental Files for "Population genomics of the facultatively asexual duckweed *Spirodela polyrhiza*"

**Table S1**. Geographic origin of 38 *S. polyrhiza* isolates sequenced. 26 isolates were collected from the wild and 12 were obtained from the Rutgers University Stock Centre (identifier RU)

| Identifier | Samples | Location | Latitude | Longitude |
|---|---|---|---|---|
| BP | 1 | Beaver Pond, Victoria, BC Canada | 48.508899 | -123.399935 |
| RB | 3 | Rithet's Bog, Victoria, BC Canada | 48.493615 | -123.373564 |
| RR | 2 | Royal Roads, Victoria, BC Canada | 48.431702 | -123.473396 |
| GP | 6 | Grenadier Pond, High Park, Toronto ON Canada | 43.642883 | -79.467660 |
| RD | 1 | Riverdale Farm, Toronto ON Canada | 43.665699 | -79.358969 |
| DD | 1 | Joker's Hill, Koffler Scientific Reserve, Toronto, ON Canada | 43.653226 | -79.383184 |
| ML | 2 | Mud Lake, Ottawa ON Canada | 45.371929 | -75.794516 |
| HF | 2 | Hennigar's Farm Market pond, Wolfville, Nova Scotia Canada | 45.085497 | -64.385739 |
| CC | 3 | Cowan Creek, Oklahoma USA | 35.359751 | -97.487380 |
| RC | 3 | Rock Creek, Oklahoma USA | 35.580741 | -97.576604 |
| RL | 2 | Johnson Lake, Raleigh North Carolina USA | 35.761433 | -78.714317 |
| RU-99 | 1 | Florida, Suwannee Co., U.S. 90 USA | 29.328850 | -83.144294 |
| RU-100 | 1 | Wisconsin, Walworth Co., Rice L.USA | 42.828954 | -88.541956 |
| RU-102 | 1 | Arkansas, Crittenden Co., Wapanocca USA | 35.373509 | -90.253840 |
| RU-103 | 1 | California, Fresno Co., Centerville USA | 36.733611 | -119.496389 |
| RU-186 | 1 | Tennessee, Obion Co., Reelfoot L. USA | 36.385680 | -89.106758 |
| RU-195 | 1 | Oregon, Douglas Co., Tahkenitch L. USA | 43.803758 | -124.125334 |
| RU-206 | 1 | Tamil Nadu, Pondicherry India | 11.919336 | 79.773631 |
| RU-408 | 1 | Gironde, Bordeaux, Marais de Blanquefort France | 44.933333 | -0.566667 |
| RU-410 | 1 | Veracruz, Coatzacoalcos Mexico | 18.134478 | -94.458986 |
| RU-412 | 1 | Texas, Davis Co., Tohyahvale USA | 30.944313 | -103.789349 |
| RU-415 | 1 | Huila, El Juncal Colombia | 2.826201 | -75.325184 |
| RU-448 | 1 | Tabasco, Villahermosa Mexico | 17.989456 | -92.947506 |

**Table S2.** Genotypic distance for pair wise comparisons of genotypes

| | | Genotype 2 | | |
|---|---|---|---|---|
| | | AA | Aa | aa |
| Genotype 1 | AA | 0 | 0.5 | 1 |
| | Aa | 0.5 | 0 | 0.5 |
| | aa | 1 | 0.5 | 1 |

’A’ and ‘a’ represents the reference and alternate allele at a site

**Table S3.** Allelic distance for pair wise comparisons of genotypes

| | | Genotype 2 | | |
|---|---|---|---|---|
| | | AA | Aa | aa |
| Genotype 1 | AA | 0 | 0.5 | 1 |
| | Aa | 0.5 | 0.5 | 0.5 |
| | aa | 1 | 0.5 | 1 |

’A’ and ‘a’ represents the reference and alternate allele at a site

**Table S4**. Heterozygosity and within/between group genotypic distance for the 12 *S. polyrhiza* genets

| Samples in genet | No. samples | Heterozygosity | Within genet | | Between genets | | No. unique alt sites |
|---|---|---|---|---|---|---|---|
| | | | mean | sd | mean | sd | |
| CC1_1, CC3_3, CC4_1, RC1_1, RC3_2, RC2_3 | 6 | 0.000595 | 1.05E-04 | 4.94E-05 | 6.04E-04 | 4.52E-04 | 10656 |
| DD7, GP14 | 2 | 0.000634 | 6.49E-06 | - | 5.64E-04 | 3.68E-04 | 7679 |
| GP10_3, GP2_3, GP4_2, GP6_5, GP8_1 | 5 | 0.000544 | 1.00E-05 | 3.51E-06 | 5.77E-04 | 3.92E-04 | 7132 |
| HFA10, HFA11 | 2 | 0.000513 | 7.14E-06 | - | 6.15E-04 | 3.40E-04 | 13635 |
| ML1_1, ML3_1 | 2 | 0.000619 | 7.48E-06 | - | 5.83E-04 | 3.85E-04 | 7547 |
| RD24 | 1 | 0.000607 | - | - | 5.58E-05 | 3.92E-04 | 5388 |
| BP1, RB1_1, RB2_7, RB3_7, RR3_6 | 5 | 0.000843 | 1.67E-05 | 4.14E-06 | 6.12E-04 | 3.92E-04 | 10350 |
| RR2_1 | 1 | 0.000878 | - | - | 4.10E-04 | 2.93E-04 | 250 |
| RL1_1, RL2_1, RU100, RU102, RU186, RU408, RU410 | 7 | 0.000526 | 6.89E-06 | 1.78E-07 | 6.32E-04 | 4.88E-04 | 8930 |
| RU448, RU99 | 2 | 0.001854 | 1.95E-05 | - | 2.00E-03 | 4.73E-04 | 223652 |
| RU103, RU206 | 2 | 0.000198 | 4.50E-06 | - | 6.35E-04 | 4.12E-04 | 2182 |
| RU195 | 1 | 0.000512 | - | - | 5.70E-04 | 4.11E-04 | 9215 |

*'No. unique alt sites' represents the number of variant sites where only the focal genotype possesses the alternate allele (i.e. all other genotypes are homozygous for the reference allele).

**Table S5**. Observed heterozygosity for 36 *S. polyrhiza* samples

| Sample | Heterozygosity |
|---|---|
| BP1 | 0.000762448 |
| CC1_1 | 0.000577312 |
| CC3_3 | 0.000564202 |
| CC4_1 | 0.000589003 |
| DD7 | 0.000604891 |
| GP10_3 | 0.00051489 |
| GP14_4 | 0.000603399 |
| GP2_3 | 0.00050829 |
| GP4_2 | 0.000484295 |
| GP6_5 | 0.000555587 |
| GP8_1 | 0.000527574 |
| HFA10 | 0.00048876 |
| HFB11 | 0.000487266 |
| ML1_1 | 0.000594203 |
| ML3_1 | 0.000601359 |
| RB1_1 | 0.000796753 |
| RB2_7 | 0.00077776 |
| RB3_7 | 0.000790806 |
| RC1_1 | 0.000571206 |
| RC3_2 | 0.000554118 |
| RC_2_3 | 0.000627124 |
| RD24 | 0.000607138 |
| RL1_1 | 0.000477178 |
| RL2_1 | 0.000491936 |
| RR2_1 | 0.000878052 |
| RR3_6 | 0.00081211 |
| RU100 | 0.000510088 |
| RU102 | 0.000516237 |
| RU103 | 0.00019054 |
| RU186 | 0.000510061 |
| RU195 | 0.000511739 |
| RU206 | 0.000193301 |
| RU408 | 0.000489469 |
| RU410 | 0.000518707 |
| RU448 | 0.001785724 |
| RU99 | 0.001806105 |

**Table S6**. Diversity statistics for 11 *S. polyrhiza* genets.

| | $\theta_w$ | $\pi$ | Tajima's D |
|---|---|---|---|
| All sites | | | |
| 0fold | 0.000253 | 0.000229 | -0.402 |
| 4fold | 0.000479 | 0.000463 | -0.143 |
| 2 or 3 fold | 0.000371 | 0.000351 | -0.229 |
| Intergenic | 0.000747 | 0.000732 | -0.0826 |
| Intronic | 0.000460 | 0.000449 | -0.107 |
| 100bp upstream | 0.000753 | 0.000738 | -0.0850 |
| Low expression | | | |
| 0fold | 0.000328 | 0.000306 | -0.275 |
| 4fold | 0.000543 | 0.000549 | 0.0440 |
| 2 or 3 fold | 0.000426 | 0.000402 | -0.239 |
| Intergenic | 0.000747 | 0.000732 | -0.0826 |
| Intronic | 0.000547 | 0.000531 | -0.122 |
| 100bp upstream | 0.000753 | 0.000738 | -0.0850 |
| Med-low expression | | | |
| 0fold | 0.000237 | 0.000212 | -0.438 |
| 4fold | 0.000441 | 0.000420 | -0.196 |
| 2 or 3 fold | 0.000334 | 0.000326 | -0.0994 |
| Intergenic | 0.000753 | 0.000738 | -0.0849 |
| Intronic | 0.000431 | 0.000424 | -0.0719 |
| 100bp upstream | 0.000546 | 0.000549 | 0.0240 |

| | | | |
|---|---|---|---|
| Med-high expression | | | |
| 0fold | 0.000203 | 0.000181 | -0.466 |
| 4fold | 0.000422 | 0.000398 | -0.240 |
| 2 or 3 fold | 0.000322 | 0.000302 | -0.263 |
| Intergenic | 0.000753 | 0.000738 | -0.0849 |
| Intronic | 0.000411 | 0.000401 | -0.101 |
| 100bp upstream | 0.000546 | 0.000549 | 0.0240 |
| High expression | | | |
| 0fold | 0.000181 | 0.000154 | -0.608 |
| 4fold | 0.000470 | 0.000445 | -0.228 |
| 2 or 3 fold | 0.000348 | 0.000320 | -0.334 |
| Intergenic | 0.000753 | 0.000738 | -0.0850 |
| Intronic | 0.000430 | 0.000417 | -0.126 |
| 100bp upstream | 0.000546 | 0.000549 | 0.0240 |
| No expression | | | |
| 0fold | 0.000762 | 0.000722 | -0.222 |
| 4fold | 0.000894 | 0.000878 | -0.0711 |
| 2 or 3 fold | 0.000934 | 0.000873 | -0.273 |
| Intergenic | 0.000753 | 0.000738 | -0.0850 |
| Intronic | 0.000104 | 0.00100 | -0.155 |
| 100bp upstream | 0.000546 | 0.000550 | 0.0278 |

**Table S7**. Diversity statistics for 11 *S.polyrhiza* genets for low expressed genes separated by level of sequence similarity to *Sorghum bicolor*, *Zea mays*, *Oryza sativa*.

| | $\theta_w$ | $\pi$ | Tajima's D |
|---|---|---|---|
| Low similarity | | | |
| 0fold | 0.000452429522958 | 0.000434982872854 | -0.161096535746 |
| 4fold | 0.000699153392111 | 0.000635904869659 | -0.37527036932 |
| 2 or 3 fold | 0.00052392749442 | 0.000505620140483 | -0.144643653089 |
| Intergenic | 0.000752803273269 | 0.000737576671008 | -0.0849079204022 |
| Intronic | 0.00071623660709 | 0.00069698067641 | -0.112757418504 |
| 100bp upstream | 0.000546076559768 | 0.000549422307173 | 0.0256872477884 |
| Medium similarity | | | |
| 0fold | 0.000320787109443 | 0.000305869410878 | -0.194526665512 |
| 4fold | 0.000543999250826 | 0.000557815939786 | 0.105767184666 |
| 2 or 3 fold | 0.000405443665052 | 0.000381234647445 | -0.248340163601 |
| Intergenic | 0.000752803273269 | 0.000737576671008 | -0.0849079204022 |
| Intronic | 0.000464319281199 | 0.000459767519553 | -0.0411032191338 |
| 100bp upstream | 0.000546076559768 | 0.000549422307173 | 0.0256872477884 |
| High similarity | | | |
| 0fold | 0.000289306037445 | 0.000261308377676 | -0.405332614955 |
| 4fold | 0.000488802808503 | 0.000512502009607 | 0.202461623273 |
| 2 or 3 fold | 0.000400918458592 | 0.000375234870958 | -0.267465943706 |
| Intergenic | 0.000752803273269 | 0.000737576671008 | -0.0849079204022 |
| Intronic | 0.000478673802757 | 0.000452218385782 | -0.231821146739 |
| 100bp upstream | 0.000546076559768 | 0.000549422307173 | 0.0256872477884 |

**Table S8**. Diversity statistics for 11 *S.polyrhiza* genets for medium-low expressed genes separated by level of sequence similarity to *Sorghum bicolor*, *Zea mays*, *Oryza sativa*.

| | $\theta_w$ | $\pi$ | Tajima's D |
|---|---|---|---|
| Low similarity | | | |
| 0fold | 0.000324246520039 | 0.00031849564278 | -0.0738342780917 |
| 4fold | 0.000518151570512 | 0.000510157486936 | -0.0634713617366 |
| 2 or 3 fold | 0.000408459326195 | 0.000454714632628 | 0.465240671245 |
| Intergenic | 0.000752812676418 | 0.000737592684983 | -0.0848699964611 |
| Intronic | 0.00048989495121 | 0.000499379313603 | 0.0811359544886 |
| 100bp upstream | 0.000545757707525 | 0.000548879660074 | 0.0239830300589 |
| Medium similarity | | | |
| 0fold | 0.000255463930957 | 0.000235264620463 | -0.330725826324 |
| 4fold | 0.000431345049715 | 0.000399962603197 | -0.302814266956 |
| 2 or 3 fold | 0.000358579904496 | 0.000351235639829 | -0.0852712534539 |
| Intergenic | 0.000752812676418 | 0.000737592684983 | -0.0848699964611 |
| Intronic | 0.000438640059418 | 0.000434164461988 | -0.0428081673863 |
| 100bp upstream | 0.000545757707525 | 0.000548879660074 | 0.0239830300589 |
| High similarity | | | |
| 0fold | 0.000212324128942 | 0.000181604219648 | -0.606074287576 |
| 4fold | 0.000429950980143 | 0.000415099476086 | -0.144383110195 |
| 2 or 3 fold | 0.000311960527841 | 0.000295712967559 | -0.217619919104 |
| Intergenic | 0.000752812676418 | 0.000737592684983 | -0.0848699964611 |
| Intronic | 0.000416681514367 | 0.000404802613261 | -0.11963729621 |
| 100bp upstream | 0.000545757707525 | 0.000548879660074 | 0.0239830300589 |

**Table S9**. Diversity statistics for 11 *S. polyrhiza* genets for medium-high expressed genes separated by level of sequence similarity to *Sorghum bicolor*, *Zea mays*, *Oryza sativa*.

| | $\theta_w$ | $\pi$ | Tajima's D |
|---|---|---|---|
| Low similarity | | | |
| 0fold | 0.000272596802361 | 0.000263443329172 | -0.139275602345 |
| 4fold | 0.000365865174662 | 0.000301579850896 | -0.712695912831 |
| 2 or 3 fold | 0.000280143726364 | 0.000288822152995 | 0.125208456414 |
| Intergenic | 0.000752813055148 | 0.000737593056056 | -0.0848699964611 |
| Intronic | 0.000433097929773 | 0.000425904063495 | -0.0696134621662 |
| 100bp upstream | 0.000545748411207 | 0.000548870310577 | 0.0239830300589 |
| Medium similarity | | | |
| 0fold | 0.000252224237486 | 0.00023742599098 | -0.245195243704 |
| 4fold | 0.000452792670482 | 0.000435608528012 | -0.157745203136 |
| 2 or 3 fold | 0.000368939282488 | 0.000347907709459 | -0.236932726775 |
| Intergenic | 0.000752813055148 | 0.000737593056056 | -0.0848699964611 |
| Intronic | 0.000417271217839 | 0.000410598247594 | -0.0670923452339 |
| 100bp upstream | 0.000545748411207 | 0.000548870310577 | 0.0239830300589 |
| High similarity | | | |
| 0fold | 0.000175825433372 | 0.000148222135127 | -0.657443774403 |
| 4fold | 0.000417782622587 | 0.000396085946346 | -0.217097191452 |
| 2 or 3 fold | 0.000309497597787 | 0.000284927547964 | -0.331785198173 |
| Intergenic | 0.000752813055148 | 0.000737593056056 | -0.0848699964611 |
| Intronic | 0.000405763304278 | 0.000394630910854 | -0.11514367613 |
| 100bp upstream | 0.000545748411207 | 0.000548870310577 | 0.0239830300589 |

**Table S10**. Diversity statistics for 11 *S.polyrhiza* genets for high expressed genes separated by level of sequence similarity to *Sorghum bicolor*, *Zea mays*, *Oryza sativa*.

| | $\theta_w$ | $\pi$ | Tajima's D |
|---|---|---|---|
| Low similarity | | | |
| 0fold | 0.000266198317664 | 0.000249067939485 | -0.26618272214 |
| 4fold | 0.000354445968287 | 0.000330386854415 | -0.273154696725 |
| 2 or 3 fold | 0.000365439545134 | 0.000363470529432 | -0.0218306606949 |
| Intergenic | 0.000752812373434 | 0.000737592388125 | -0.0848699964611 |
| Intronic | 0.000444636472758 | 0.000422869928109 | -0.20538771893 |
| 100bp upstream | 0.000545765144808 | 0.000548887139901 | 0.0239830300589 |
| Medium similarity | | | |
| 0fold | 0.00022643361714 | 0.000199603842783 | -0.494525542641 |
| 4fold | 0.000472980128563 | 0.000459695533942 | -0.116603355252 |
| 2 or 3 fold | 0.000340874848807 | 0.00032971041465 | -0.135753091711 |
| Intergenic | 0.000752812373434 | 0.000737592388125 | -0.0848699964611 |
| Intronic | 0.000444636472758 | 0.000422869928109 | -0.20538771893 |
| 100bp upstream | 0.000545765144808 | 0.000548887139901 | 0.0239830300589 |
| High similarity | | | |
| 0fold | 0.000145210544633 | 0.000117255488944 | -0.804694001273 |
| 4fold | 0.000487975472987 | 0.00045671778503 | -0.267493056279 |
| 2 or 3 fold | 0.000350229284104 | 0.000310454784616 | -0.47394846962 |
| Intergenic | 0.000752812373434 | 0.000737592388125 | -0.0848699964611 |
| Intronic | 0.000418672941106 | 0.000409421391892 | -0.0927278880616 |
| 100bp upstream | 0.000545765144808 | 0.000548887139901 | 0.0239830300589 |

**Table S11**. Coefficients from linear model of diversity ($\pi$) of different site types (per 50 kb window) as function of coding site (CDS) density, % GC, and the CpG/GpC ratio. Required that all 50 kb windows had at least SNP of each site type (intergenic, intronic, 0-fold and 4-fold).

| Site type | Variable | Estimate | t | P-value |
|---|---|---|---|---|
| Intergenic | CDS density | -0.00086 | -2.48 | 0.013 |
| | GC content | -0.00099 | -2.37 | 0.018 |
| | CpG/GpC ratio | -0.00062 | -3.69 | < 0.0001 |
| Intronic | CDS density | -0.00119 | -4.64 | < 0.0001 |
| | GC content | 0.00001 | 0.03 | 0.977 |
| | CpG/GpC ratio | -0.00028 | -2.26 | 0.024 |
| 0-fold | CDS density | -0.00151 | -6.42 | < 0.0001 |
| | GC content | 0.00015 | 0.54 | 0.593 |
| | CpG/GpC ratio | -0.00013 | -1.18 | 0.239 |
| 4-fold | CDS density | -0.00249 | -6.03 | < 0.0001 |
| | GC content | -0.00045 | -0.90 | 0.367 |
| | CpG/GpC ratio | 0.00005 | 0.27 | 0.789 |

**Table S12.** List of 131 putative hotspots identified among 11 *S. polyrhiza* genets.

| Scaffold | Start position (kb) | Length (kb) |
|---|---|---|
| 1 | 637 | 2 |
| 1 | 2567 | 3 |
| 1 | 3495 | 2 |
| 1 | 5548 | 2 |
| 1 | 6450 | 5 |
| 1 | 6547 | 3 |
| 1 | 7536 | 1 |
| 1 | 7540 | 2 |
| 1 | 7547 | 4 |
| 2 | 367 | 2 |
| 2 | 614 | 1 |
| 2 | 622 | 1 |
| 2 | 722 | 3 |
| 2 | 751 | 4 |
| 2 | 1795 | 1 |
| 2 | 3901 | 4 |
| 2 | 5278 | 2 |
| 3 | 695 | 5 |
| 3 | 814 | 1 |
| 3 | 851 | 4 |
| 3 | 1066 | 1 |
| 3 | 1470 | 2 |
| 3 | 1503 | 4 |
| 3 | 4824 | 4 |
| 3 | 5203 | 2 |
| 3 | 6041 | 3 |
| 3 | 6463 | 1 |
| 3 | 7065 | 1 |
| 3 | 7066 | 1 |
| 4 | 2299 | 1 |
| 4 | 3690 | 3 |
| 4 | 3707 | 1 |
| 4 | 4685 | 1 |
| 4 | 7177 | 3 |
| 5 | 3793 | 2 |
| 5 | 4248 | 3 |
| 5 | 4418 | 2 |
| 5 | 4597 | 3 |
| 5 | 4618 | 2 |
| 5 | 6002 | 2 |
| 6 | 964 | 2 |
| 6 | 2987 | 4 |
| 6 | 4910 | 2 |
| 6 | 5573 | 4 |

| | | |
|---|---|---|
| 7 | 1473 | 2 |
| 7 | 3038 | 1 |
| 7 | 4602 | 3 |
| 7 | 4607 | 2 |
| 7 | 6232 | 3 |
| 8 | 1465 | 1 |
| 8 | 1569 | 1 |
| 8 | 2578 | 2 |
| 9 | 2245 | 2 |
| 9 | 2528 | 2 |
| 9 | 3229 | 3 |
| 10 | 1038 | 1 |
| 10 | 1611 | 5 |
| 10 | 2030 | 2 |
| 10 | 2089 | 4 |
| 10 | 3162 | 2 |
| 10 | 3407 | 3 |
| 10 | 3503 | 3 |
| 11 | 303 | 4 |
| 11 | 1453 | 2 |
| 11 | 1521 | 2 |
| 11 | 2216 | 4 |
| 11 | 2355 | 2 |
| 11 | 3371 | 4 |
| 11 | 3385 | 3 |
| 11 | 3527 | 5 |
| 11 | 3918 | 1 |
| 12 | 1327 | 3 |
| 12 | 2077 | 3 |
| 12 | 2149 | 3 |
| 12 | 2157 | 2 |
| 12 | 3097 | 2 |
| 13 | 2136 | 1 |
| 13 | 4023 | 1 |
| 14 | 1293 | 1 |
| 14 | 1394 | 1 |
| 14 | 1870 | 1 |
| 14 | 1937 | 2 |
| 14 | 2024 | 2 |
| 14 | 2060 | 6 |
| 14 | 2341 | 2 |
| 14 | 2953 | 2 |
| 14 | 3655 | 2 |
| 14 | 3695 | 1 |
| 15 | 1109 | 6 |
| 15 | 3019 | 2 |
| 15 | 3182 | 1 |

| | | |
|---|---|---|
| 15 | 3193 | 2 |
| 16 | 915 | 1 |
| 17 | 52 | 1 |
| 17 | 1303 | 3 |
| 17 | 1325 | 6 |
| 18 | 2740 | 4 |
| 18 | 2826 | 3 |
| 18 | 2836 | 1 |
| 19 | 2 | 2 |
| 21 | 665 | 1 |
| 21 | 2942 | 2 |
| 22 | 1540 | 1 |
| 23 | 1053 | 3 |
| 23 | 2279 | 3 |
| 24 | 1080 | 3 |
| 24 | 1656 | 2 |
| 25 | 583 | 2 |
| 25 | 1817 | 3 |
| 25 | 1840 | 2 |
| 25 | 2027 | 4 |
| 25 | 2084 | 2 |
| 26 | 289 | 3 |
| 26 | 304 | 1 |
| 26 | 410 | 4 |
| 26 | 1064 | 3 |
| 26 | 1228 | 3 |
| 26 | 1245 | 4 |
| 26 | 1770 | 3 |
| 27 | 862 | 2 |
| 27 | 914 | 2 |
| 27 | 923 | 2 |
| 27 | 1627 | 3 |
| 29 | 8 | 3 |
| 30 | 537 | 3 |
| 30 | 616 | 1 |
| 30 | 1003 | 2 |
| 31 | 212 | 3 |
| 31 | 420 | 2 |
| 31 | 642 | 2 |
| 32 | 918 | 2 |

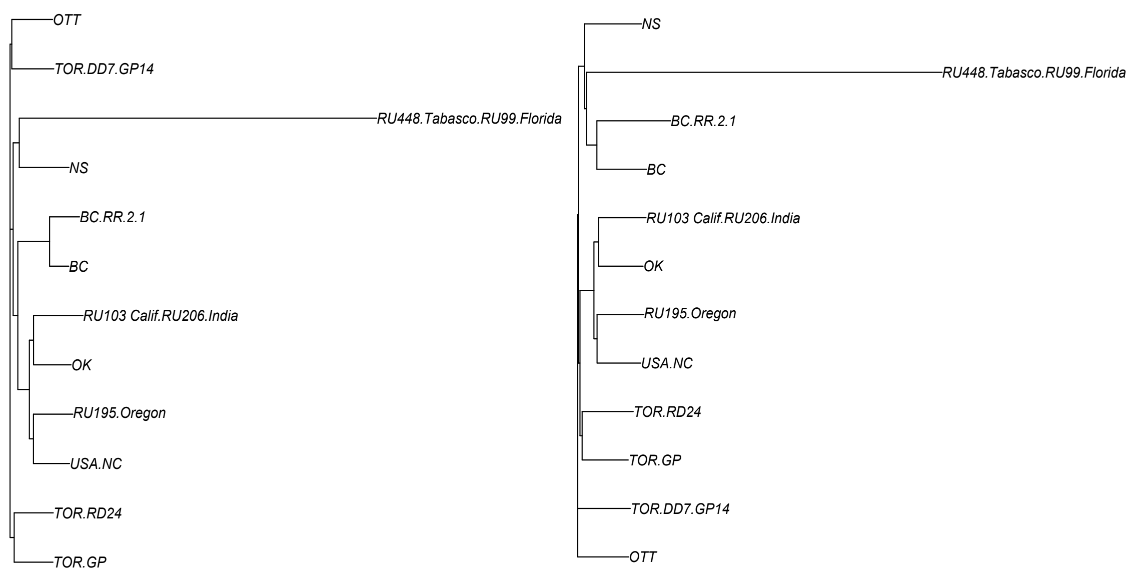


**Figure S1**. Neighbour joining tree for the 12 *S. polyrhiza* genets constructed based on pairwise genotypic distance (left) and allelic distance (right).
